# Frequency variation and dose modification of benznidazole administration for the treatment of *Trypanosoma cruzi* infection in mice, dogs and non-human primates

**DOI:** 10.1101/2023.02.01.526739

**Authors:** Juan M. Bustamante, Brooke E. White, Gregory K. Wilkerson, Carolyn L. Hodo, Lisa D. Auckland, Wei Wang, Stephanie McCain, Sarah A. Hamer, Ashley B. Saunders, Rick L. Tarleton

## Abstract

*Trypanosoma cruzi* naturally infects a broad range of mammalian species and frequently results in the pathology that has been most extensively characterized in human Chagas disease. Currently employed treatment regimens fail to achieve parasitological cure of *T. cruzi* infection in the majority of cases. In this study, we have extended our previous investigations of more effective, higher dose, intermittent administration protocols using the FDA-approved drug benznidazole (BNZ), in experimentally infected mice and in naturally infected dogs and non-human primates (NHP). Collectively these studies demonstrate that twice-weekly administration of BNZ for more than 4 months at doses that are ∼2.5-fold that of previously used daily dosing protocols, provided the best chance to obtain parasitological cure. Dosing less frequently or for shorter time periods was less dependable in all species. Prior treatment using an ineffective dosing regimen in NHPs did not prevent the attainment of parasitological cure with an intensified BNZ dosing protocol. Furthermore, parasites isolated after a failed BNZ treatment showed nearly identical susceptibility to BNZ as those obtained prior to treatment, confirming the low risk of induction of drug resistance with BNZ and the ability to adjust the treatment protocol when an initial regimen fails. These results provide guidance for the use of BNZ as an effective treatment for *T. cruzi* infection and encourage its wider use, minimally in high value dogs and at-risk NHP, but also potentially in humans, until better options are available.

## Introduction

Chagas disease is a vector-borne parasitic disease caused by the hemoflagellate protozoan *Trypanosoma cruzi* and is one of the major world-wide causes of infection-induced myocardial disease. *T. cruzi* is widely distributed and transmitted to a broad range of mammals, from the southern United States to the far south of Chile and Argentina. Managing Chagas disease is particularly challenging; human infections are not routinely diagnosed before health-impacting tissue damage is substantial and without treatment, the infection generally persists for life. Systematic vector control can limit new human infections but can be expensive and labor intensive and may have less impact on the rate of new infections in animals in some settings [1-3]. The current treatment for human *T. cruzi* infection depends on two nitroheterocyclic compounds, benznidazole (BNZ) and nifurtimox (NFX), that are given in an intensive treatment regimen (usually 30-60 twice-daily doses for BNZ and 30-120 daily doses for NFX). These treatment protocols are frequently accompanied by adverse events, and in addition have unpredictable efficacy [4]. Collectively, these attributes result in a low use rate and in a relatively high degree of treatment interruption when used [5].

Despite these significant limitations, BNZ’s anti-*T. cruzi* activity is particularly interesting given that a single dose administered *in vivo* can rapidly decrease parasite load by 90% or more [6] and abbreviated treatment protocols can achieve parasitological cure in mice and humans [5, 7, 8]. Nevertheless, the conventional 30-day and even 60-day treatment regimens have failure rates exceeding 50% [9]. These discrepancies in treatment outcomes are likely due to a complicated set of factors that include the initial infecting dose and the genetics of the infecting paarasites, the length of infection and the host immune response to the infection, among others. We have also recently reported that not all parasites in a host at any one time are equally susceptible the trypanocidal effects of drugs such as BNZ, a result of the ability of some *T. cruzi* amastigotes to enter a transient state of dormancy [6, 8].

Given that the availability of new drugs for treating *T. cruzi* infection are likely many years away, a primary goal of this study was to determine if dose modification of an existing drug could improve treatment outcomes. For this purpose, we continued our work in experimentally infected mice, where highly controlled and rigorously assessed studies can be conducted. Taking advantage of the incidence of *T. cruzi* infection in non-human primates in U.S. research and zoological facilities, and very high incidence and high impact infection in working dogs in the southern U.S. [3, 10, 11], we further explored these treatment modifications in these hosts with the advantage of variable host and parasite genetics, infection dose and infection length that is also a property of infections in humans. The results, while not identifying a single failsafe treatment protocol, nevertheless provide guidance on the more effective use of BNZ in multiple species, possibly including humans.

## Results

### A biweekly treatment with benznidazole can cure mice infected with *T. cruzi*

We previously demonstrated that high dose BNZ treatments given once per week for 30 weeks (but not 20 weeks) achieved parasitological cure in mice chronically infected with *T. cruzi* [8]. Further adjusting this intermittent protocol in anticipation of its potential use in the treatment of chronic infections in the field and with the goal of reducing the overall treatment period, we tested the impact of dosing twice-per-week BNZ at 250mg/kg (2.5X that of the standard daily dose for mice) for 8 weeks. The period of 8 weeks was selected based on strong evidence of clearance of both active and dormant amastigotes from the tissues of mice with acute *T. cruzi* infections using this treatment period [8]. For these studies, we once again used mice chronically infected (480 days) with the Colombiana strain of *T. cruzi* that is relatively difficult to clear *in vivo* [7, 8].

Mice with a persistent *T. cruzi* infection maintain a relatively low frequency of CD127-expressing (CD127hi) CD8+ T cells specific for the TSKb20 *T. cruzi* trans-sialidase (ts)-derived epitope while mice with reduced parasite load or cured infections have an increased proportion of *T. cruzi*-specific, CD127hi CD8+ T cells [7, 8, 12]. This marker of reduced antigen exposure was observed during treatment (6 weeks) and 2 weeks after the end of treatment (Fig. 1B) and complete parasite clearance was confirmed by the absence of a detectable PCR signal in multiple tissues (Fig 1C) and by negative hemocultures (Fig. 1D).

**Figure 1.**
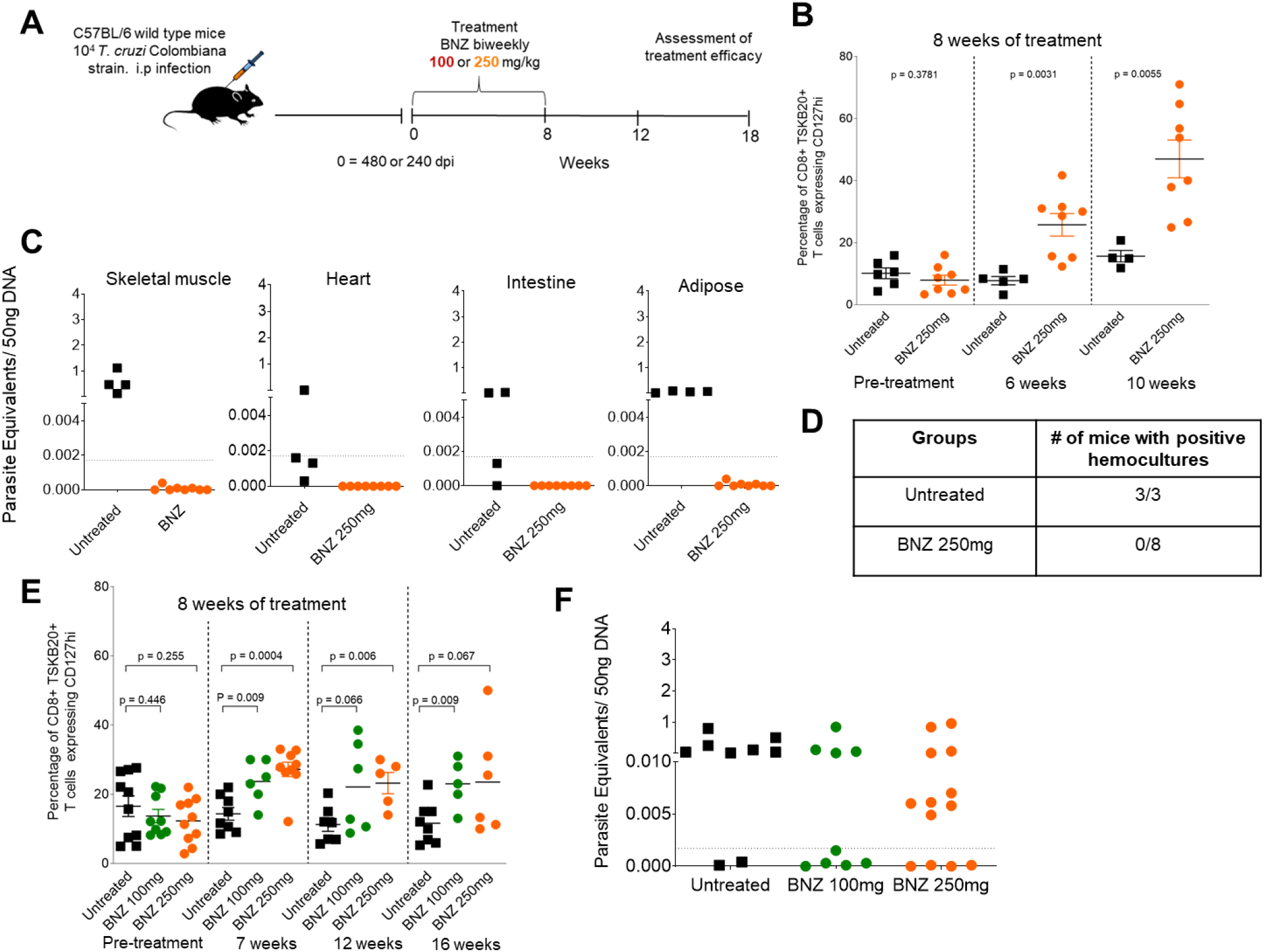
**(A)** Schematic representation of infection, treatment and assessment of treatment efficacy. C57BL/6 wild-type mice (6-13 mice in each group) were infected intraperitoneally with 10^4^ trypomastigotes of the Colombiana strain of *T. cruzi* and left untreated or treated biweekly, starting at 480 days (Exp 1 panels B-D) or 240 days (Exp 2 panels E and F) post-infection, with BNZ at 100 or 250mg/kg concentration over 8 weeks. **(B)** Expression of the T_cm_ marker CD127 in blood on CD8^+^ TSKB20-tetramer+ T cells from individual mice untreated or treated at 12 weeks. **(C)** *T. cruzi* DNA isolated from tissues of untreated or treated mice at 12 weeks post-initiation of treatment and assayed by qPCR. A statistically significant difference was set at p=0.05. **(D)** Blood from these mice collected at 12 weeks was submitted to hemoculture assays and analyzed for parasite growth for >60 days. Cultures were negative for parasites in the BNZ treated group and positive for the untreated ones. **(E)** Expression of the T_cm_ marker CD127 in blood on CD8^+^ TSKB20-tetramer+ T cells from individual mice infected as in **(A)** for 240 days and left untreated or treated biweekly with BNZ at 100 or 250mg/kg over 8 weeks. **(F)** qPCR detection of *T. cruzi* DNA in skeletal muscle tissue from untreated or treated mice at 18 weeks post-initiation of treatment.

We repeated this experiment with 3 modifications: 1) mice were infected for 240 days at the time of treatment, 2) inclusion of a lower BNZ dose group (100 mg/kg; the standard daily dose but given twice per week), and 3) terminating the experiment for collection of tissues at 10 weeks after the end of treatment, rather than at 2 weeks. Although the frequency of *T. cruzi*-specific CD8+ T cell expressing CD127 (a surrogate indicator of reduced parasite load) increased in both the treated groups, this indicator was not consistently increased throughout the treatment period, as observed in the first experiment (Fig. 1E). Furthermore, unlike the first experiment, skeletal muscle tissue from the majority of mice in the high dose -treated group remained *T. cruzi* DNA PCR-positive at 10 weeks after the end of the 8-week treatment period (Fig. 1F). Collectively, these results demonstrate that while twice weekly, high dose BNZ treatment can cure *T. cruzi* in as few as 8 weeks (Fig. 1), this effect is not consistent across different experimental groups infected for different periods of time, even using the same infecting *T. cruzi* strain.

### Chronically infected dogs treated with weekly and twice weekly benznidazole regimen

Dogs are significant hosts for *T. cruzi* infection and in certain regions of the U.S. are under intense infection pressure, with 20-30% incidence of new infections per year [3, 10]. Standard, twice daily BNZ dosing regimens in dogs with experimental *T. cruzi* infections has a very poor efficacy record [13, 14]. However, given our relative success with higher BNZ dosing regimens in mice combined with the challenge of dosing client-owned dogs, we attempted a small trial using once per week dosing of BNZ at 20 mg/kg, which is 2-2.5X the standard daily dose. We also chose to treat for a minimum of 6 months using this regimen. Thus, these dogs received approximately the same total cumulative amount of BNZ as in standard, 60 consecutive daily dosing protocols.

All 7 dogs in the trial were *T. cruzi* seropositive by multiple tests and 5 of the 7 were blood PCR positive for *T. cruzi* during the 2 yrs prior to the start of treatment (Fig. 2A). *T. cruzi* was also detected by hemoculture in one of the 7 dogs. PCR typing indicated that 3 of the dogs had *T. cruzi* discrete typing unit (DTU) IV parasites and one had DTU I (Supplemental Table 1). The beginning of treatment in these dogs was staggered in order to assess the potential neurological impact of higher than standard BNZ dose, as previously reported in some canines under very high dose BNZ treatment [15] (see Materials and Methods). No changes in mental status or locomotory function were noted during the 27-45 week treatment period. One dog exhibited transient hair loss between weeks 10 and 12 of treatment but new hair growth was evident by week 18.

**Figure 2.**
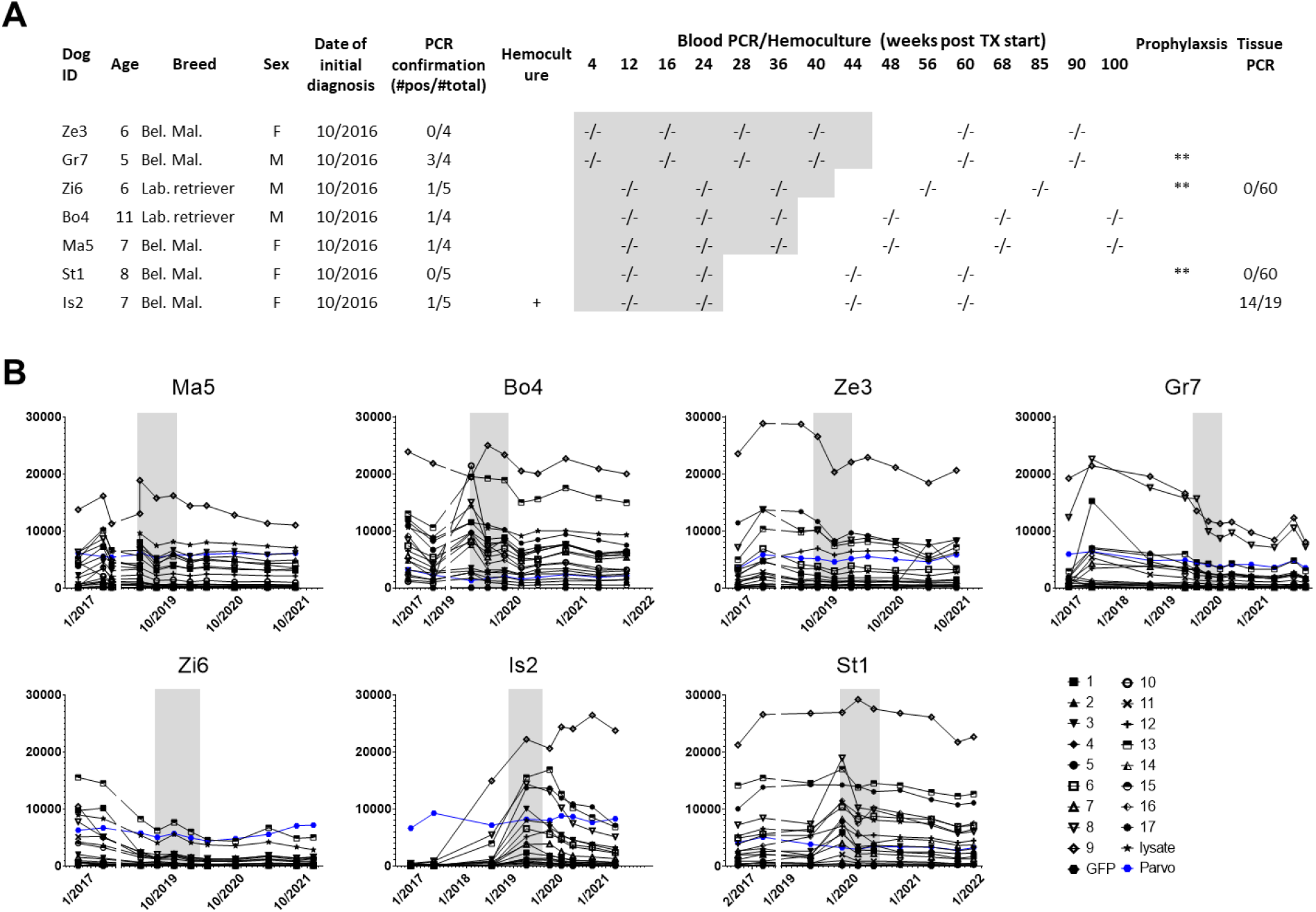
**(A)** Dog profiles and blood and tissue PCR and hemoculture results before, during (grayed region) and after treatment. At approximately 1 yr after the end of treatment, some dogs (marked with **) were placed on a proposed prophylactic regimen designed to reduce the occurrence of reinfections [10]. Tissue PCR data indicate the number of *T. cruzi* DNA-positive reactions per total number of tissue specimens tested singly or in pools of 5 from heart and skeletal muscle. **(B)** Profiles of serum antibodies to *T. cruzi* and control proteins pre-, during- and post-treatment. Treatment period for each dog indicated by gray box. Y axis is mean fluorescence intensity. Full study details are shown in Supplemental Table 1.

All BNZ-treated dogs were consistently negative for *T. cruzi* DNA in PCR screening of blood during and after treatment (up to ∼2 years in some cases; (Fig. 2A and supplemental Table 1)). Multiplex serological screening to assess declines in antibody levels to multiple *T. cruzi* proteins, an indicator of treatment efficacy in humans [16-19] and non-human primates [20], also showed declining antibody levels in most, but not all animals over this relatively short post-treatment observation period (Fig. 2B).

The culling and death of four of the dogs (Supplemental Table 1) provided an opportunity to screen tissue samples from three dogs for evidence of residual parasites in cardiac and skeletal muscle, sites of known *T. cruzi* persistence in chronically infected hosts. In addition to the indicated high dose curative treatment regimen, dogs Zi6 and St1 had received an additional 6-month, low dose BNZ prophylactic regimen intended to prevent re-infection [10], ending approximately 1 yr prior to their necropsy. A total of 30 individual samples and 6 pooled samples from 5 sites each in heart and skeletal muscle, for a total of 60 individual sites from dogs ZI6 and ST1, were negative for *T. cruzi* DNA. However, animal IS2, in which the cause of death was heartworms, was PCR positive in multiple samples from both skeletal muscle and heart (Fig. 2A). IS2 had one of the shortest treatment periods of these 7 animals (27 weeks) and also was the only animal that had increasing antibody levels to one of the multiplex antigens after the end of treatment, another indicator of failed treatment (Fig. 2B). Thus, a once per week, high dose BNZ treatment regimen had a therapeutic benefit in dogs with long-standing *T. cruzi* infection of different genetic types but failed to provide uniform cure when administered for less than 30 weeks.

### Impact of weekly benznidazole treatment in macaques with naturally acquired *T. cruzi* infection

In previous studies in cynomolgus macaques naturally infected with *T. cruzi*, we found that BNZ treatment given at the standard dose on a daily schedule of 60 consecutive days, cured <50% of the infected animals (Padilla, unpublished). This is in line with the success rate for BNZ in humans [21]. To determine if a high dose but intermittent BNZ treatment protocol such as those evaluated in mice and dogs above might be also be efficacious in chronically infected primates, we initiated a pilot study in 10 rhesus macaques with naturally acquired *T. cruzi* infection, all of which were blood PCR+ at multiple sampling times over several years (Supplemental Table 2). We initially treated 8 macaques with 37.5 mg/kg BNZ (2.5 times the standard daily dose), once per week, with the intention to continue this protocol for a period 1 year (Figure 3). However, due to COVID-19 research restrictions, treatment had to be suspended at week 28 (Fig. 3A). At this point, 5 of the 8 treated macaques were blood PCR and hemoculture negative, while 3 of the treated group remained blood PCR positive, indicating failures for this treatment protocol (Fig.3B).

**Figure 3.**
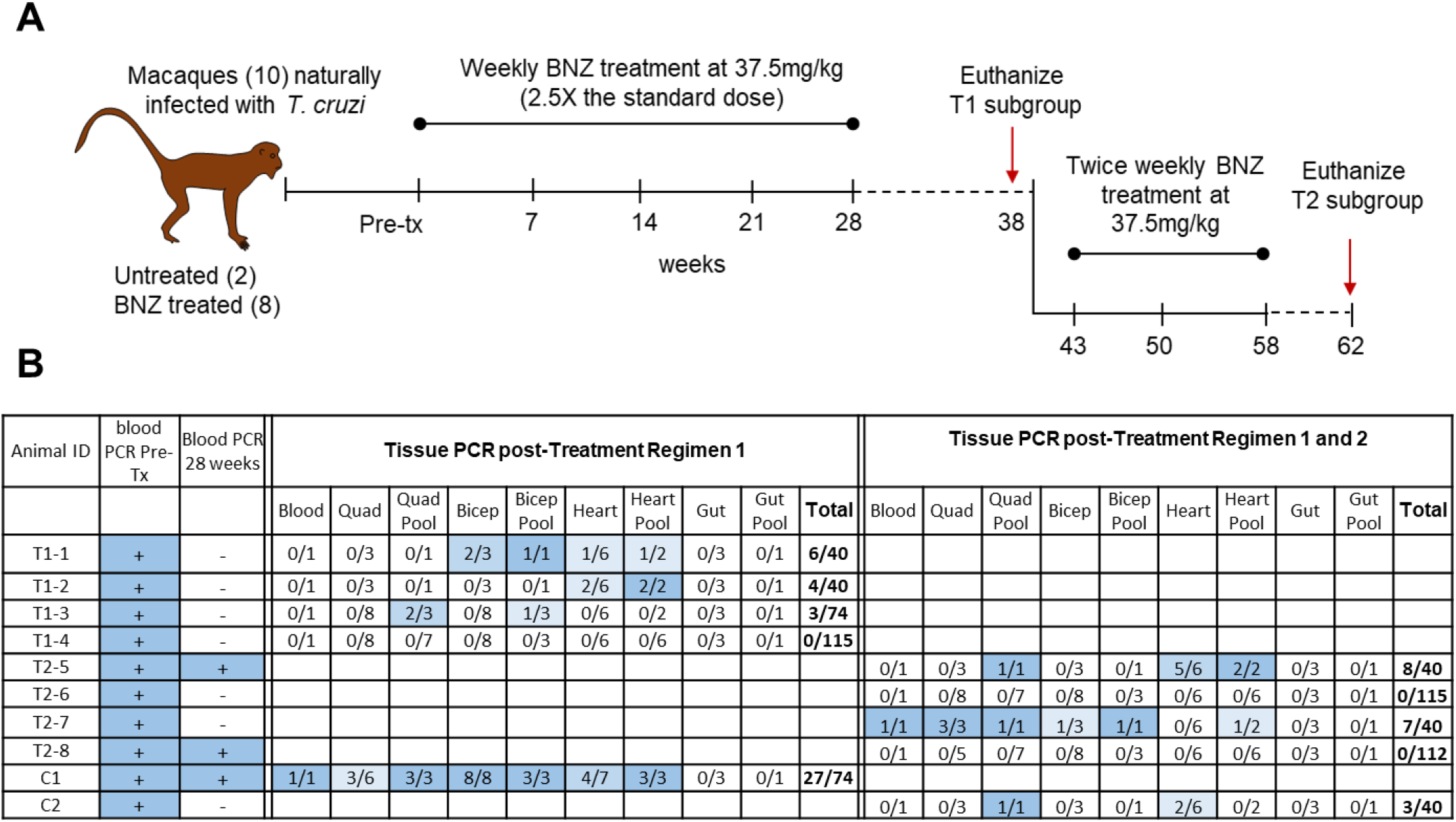
**(A)** Schematic representation of the study in chronically infected macaques treated weekly and biweekly with BNZ. A total of 10 macaques chronically infected with *T. cruzi* were used in this study, 8 were treated with BNZ weekly at a concentration of 37.5mg/kg (2.5 times the standard daily dose) over 28 weeks, follow by a cessation of the treatment between week 28 and 38 due to COVID-19 research restrictions. Two animals were kept as untreated controls. At week 38, five macaques (4 treated and 1 untreated) were necropsied to collect the tissues for assessment of parasites persistence by qPCR. With the remaining 5 macaques, one of them was continued untreated and the other 4 were treated with a second round of BNZ delivered biweekly at the same concentration as before, over a 15-week period (from week 43 to 58). At the 62-week time point this second set of macaques (4 treated and 1 untreated) were necropsied to collect the tissues for assessment of parasites persistence by qPCR. **(B)** Macaque identification, pre-treatment and end of treatment course 1 (28 weeks) blood qPCR status, and tissue PCR results at necropsy are shown. Totals for tissue PCR indicate the number of *T. cruzi* DNA-positive PCR reactions per total number of tissue specimens tested singly or in pools of 5 from the indicated tissues.

In order to confirm or not parasitological cure in the blood PCR-negative treated macaques, 4 of the 6 and 1 untreated were euthanized 10 weeks after the end of their treatment protocol and their tissues assessed by PCR for *T. cruzi* DNA, as done above for dogs (Fig.3B). *T. cruzi* DNA was detected in the quadricep, bicep and heart muscle of the untreated macaque, and in the bicep and/or heart of 3 of the 4 BNZ-treated, blood PCR-negative macaques (Fig. 3B). However, the remaining macaque was tissue and blood PCR negative.

We were able to resume treatment in the remaining 4 previously treated macaques at week 43 of the protocol (these 4 macaques had not received treatment between weeks 28 and 43; Fig. 3A)). Given the less than uniform efficacy of the 1X/week protocol and our success without adverse events of the 2X/week protocol in mice and dogs, we decided to resume treatment at the same 37.5 mg/kg dose but given 2X/week. This protocol was maintained for 15 weeks (until week 58 of the study) and was followed by a 4-week post-treatment period before screening of blood and necropsy samples for parasite DNA. Two of the 4 macaques were blood and tissue PCR negative after the second treatment phase but the 2 others remained parasite positive in blood and/or multiple tissue sites (Fig.3B).

One of the eight macaques (animal T1-3) developed a generalized moderate reddening of the skin and a maculopapular rash within one hour after the animal had been administered the twenty-fourth dose (Fig.3A) of BNZ in the first treatment period (week 24). This animal also had a mild eosinophilia identified on blood work collected from the same day (Supplemental Table 5). The skin reddening and rash responded well to diphenhydramine treatment although the eosinophilia persisted through the following week. As a result of the persistent eosinophilia, the animal was not dosed with BNZ at week 25. After the animal had recovered for two full weeks, the animal was again administered high dose BNZ to determine if the reaction was associated with drug administration. The animal had a similar dermatological reaction and no further BNZ administrations were attempted in this animal. Dermatological reactions are the most frequently reported side effects in humans and have occurred in up to 50% of patients treated using standard therapeutic regimens [22]. No other physical abnormalities associated with BNZ administration were noted in the treated macaques and the mild body weight and body temperature fluctuations observed in the animals were consistent with those expected of macaques housed under long-term study conditions (Supplemental Table 3). Seven of the eight macaques treated with the high dose BNZ protocol had transient mild elevations in alanine transaminase (ALT) that typically occurred following several months of treatment. The elevated ALT values decreased for most animals during the recovery phases of the study (Supplemental Table 4). Similar ALT changes have been noted in humans treated with 5-10 mg/kg/day [23]. No other serum chemistry abnormalities and no hematological changes were consistently identified in the treated macaques (Supplemental Tables 4, 5).

For one of the two macaques remaining PCR-positive after both the 28 week,1X per week and the 15 week, 2X per week treatment, we also isolated parasites by hemoculture after these two rounds of treatment. As one concern in the use of intermittent and gapped treatment regimens as employed herein is the possibility of selecting for parasites of greater drug resistance, we compared these parasites recovered post-treatment to parasites isolated from an untreated macaque and found no difference in the 2 lines with respect to in vitro BNZ. (Figure 4).

**Figure 4.**
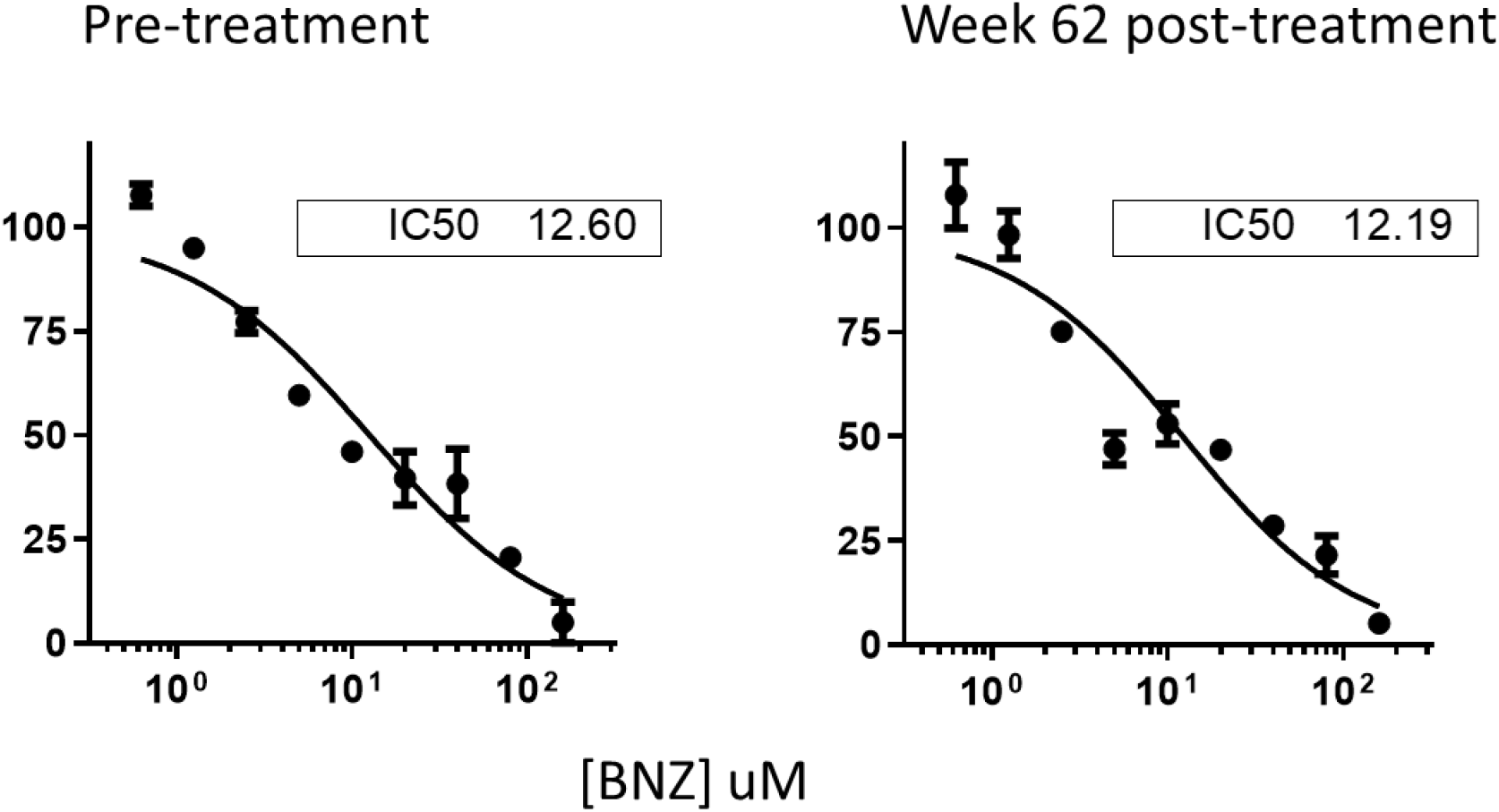
*In vitro* grown inhibition assay and calculated IC_50_ for BNZ in parasite lines isolated from hemocultures from a macaque prior to treatment (left) and from hemocultures of a macaque samples following both rounds of BNZ treatment (at week 62 of the study; right).

Lastly, the success of the above studies were of use with respect to designing treatment for a zoo-housed De Brazza’s monkey (*Cercopithecus neglectus*) with a naturally acquired *T. cruzi* infection [24]. A standard (3.5 mg/kg PO q12h x 60 days) BNZ treatment course failed to achieve clearance of *T. cruzi* in the blood (as assessed by PCR; Fig. 5). When cardiac troponin levels increased ∼16 months after this failed treatment, a higher dose (5X the standard daily dose = 35mg/kg) protocol was attempted. Weekly administration of this dose for 11 weeks also failed to achieve blood clearance of *T. cruzi*, so administration of this same dose was increased to twice weekly for 15 weeks, ultimately achieving consistent PCR negativity. As shown with respect to successfully treated dogs, declining serum antibody levels to multiple *T. cruzi* antigens supported the conclusion of a successful treatment.

**Figure 5.**
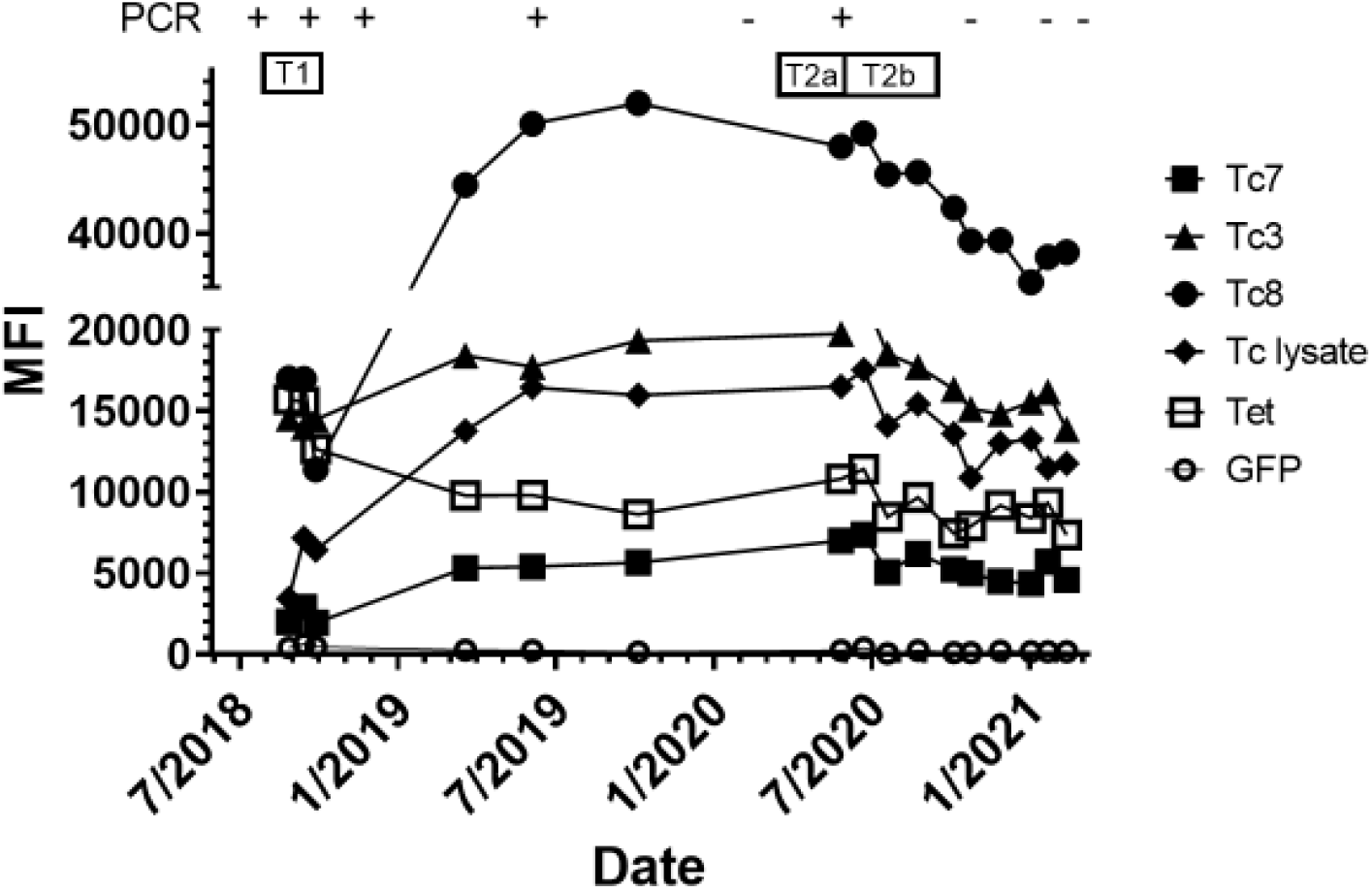
Blood PCR results and Luminex-based evaluation of anti-*T. cruzi* serum antibodies in a De Brazza’s monkey undergoing treatment with BNZ in 3 phases: T1 - 3.5 mg/kg twice daily for 60 days; T2a - 35 mg/kg once weekly for 11 weeks; and T2b - 35 mg/kg twice weekly for 15 weeks. For clarity, the graph shows only IgG profiles specific for antigens for which there was a detectable response above the level of the GFP control protein at any point during the screening period. MFI = mean fluorescence intensity

## Discussion

Although BNZ has been in use for >50 years for the treatment of *T. cruzi* infection, the drug has a reputation for a high failure rate and thus is often not employed in standard practice, despite the absence of more effective alternatives. The standard twice-daily, 30-90 day protocols currently used in humans have proven problematic in two respects: efficacy is highly variable but almost certainly well below 50% overall, and the frequency of adverse events is high, with up to 20% of subjects having to abort the protocol [22]. Our previous work has suggested that the efficacy of BNZ treatment can be significantly improved by increasing the individual dose level but administering it less frequently, so that the cumulative drug exposure is not substantially increased [7, 8].

The biological support for this high dose/intermittent BNZ treatment regimen comes from several observations. First, we have recently discovered that not all *T. cruzi* amastigotes in infected hosts are actively replicating and that these non-replicating (dormant) amastigotes are insensitive to BNZ and other anti-*T. cruzi* drugs in vitro and in vivo [6]. The longevity of these dormant forms in a state of quiescence is not known, but is likely to be highly variable, implicating that well-timed dosing over an extended period of time may be necessary to clear all parasites from a host. Second, BNZ, in addition to its potent trypanocidal activity also has modest, but significant cytostatic activity (our observations and [25]). The constant presence of BNZ achieved by the standard twice-daily dosing regimen may actually maintain a small population of amastigotes in a metabolically inactive state throughout the treatment period. Third, *T. cruzi* invades and persists in a variety of tissue types and BNZ does not reach equivalent tissue penetration in all tissue types [26].

Thus, the premise behind a high dose, intermittent treatment protocol for *T. cruzi* infection is that each dose of drug kills the majority of actively replicating parasites in all tissues, and that the interval between treatments provides time for a proportion of quiescent parasites (either naturally dormant or cytostatic drug-induced) to reinitiate metabolic activity and presumably become sensitive to the next drug dose. The optimal interval between doses as well the total period of treatment has not previously been extensively investigated. Additionally, rigorous documentation that an intermittent dosing regimen can cure naturally acquired and chronic *T. cruzi* infection has not previously been addressed.

In this study, we show in multiple species that a higher dose, weekly or twice-weekly BNZ dosing protocol has the potential to cure *T. cruzi* infection. Even though the individual drug doses are higher than in standard twice daily treatment protocols, and the period of treatment is longer, the total drug exposure is in line with existing protocols, so that the risks of adverse events may not be increased. Providing higher doses of drug at each treatment point seeks to assure that the effective concentration for killing of metabolically active parasites is achieved in all sites of parasite persistence and despite the variability in penetration of drug, and parasite residence, in these sites.

Although this study did not identify a single BNZ treatment protocol that is uniformly effective in all situations/species, it nonetheless provided important new information toward that goal. The work in four species showed that BNZ can be given at high doses, once or twice per week and over extended periods of time without toxicity and without selecting for drug resistance. Further, this work demonstrates treatment protocols that are effective in one circumstance may fail in others that look very similar. The differential outcome in the two mouse studies in this work demonstrate this. On the surface, the only variable between these experiments is the length of infection (∼400 days in exp 1 and ∼250 days in exp 2). Thus, although host and parasite genetics can and certainly do impact treatment outcomes, even when one tightly controls for those variables, the outcome is not necessarily predictable, and likely depends on other non-controllable aspects, such as the variation of parasite distribution, the frequency of dormant forms and relative potency of immune responses, all of which can affect infection outcome and drug impact. By using naturally infected dogs and NHPs rather than solely experimentally infected inbred mice as treatment recipients, this work directly encompasses these and other variables.

The impending treatment failure in the mouse experiment 2 was predictable from the pattern of CD8+ T cell memory phenotypes (Fig. 1B and E). This dependable marker of cure [7, 8, 12] was increasing in high-dose-treated mice in experiment 1 but was inconsistently elevated in experiment 2. The lack of similar “in-treatment” or post-treatment biomarkers of treatment efficacy for dogs, NHP, and humans, makes monitoring progress and assessing outcomes in these species problematic. As in many previous studies, blood PCR alone failed to detect macaques with continuing infection after treatment. Declines in anti-parasite antibody levels are highly variable between individuals and are often slow to occur and thus not useful for assessment during or soon after treatment (although they do have utility where longer-term follow-up is possible [18, 19]). Intensive screening for *T. cruzi* DNA in organ tissues provides solid evidence for cure/not-cure [20] but obviously requires termination of study animals. We took advantage of the culling of several working dogs to obtain tissue samples that solidified the evidence for parasitological cures using the high-dose, weekly BNZ protocol. Improvements in non-terminal biomarkers of treatment efficacy remains a critical need for assessing treatment outcomes in Chagas disease.

Although this work does not identify a single BNZ treatment protocol that can be expected to provide uniform cure outcomes in all cases, it strongly suggests that a high BNZ dose (minimum 2.5X the standard daily dose) administered every 3-4 days for 6 months would be a good starting point for wider evaluation. The biology of the infection supports a twice-weekly regimen as the roughly 4-5 day intracellular cycle of *T. cruzi* amastigotes would assure that few if any parasites can complete a full intracellular cycle during the interim between treatment doses, thus promoting a continuous attrition of parasites, despite the potential for dormancy. Shorter terms of treatment (< 6 months) work in some hosts but not in others – similar to the situation in humans when treatment is truncated for various reasons [27]. There is increased risk of failure with shorter treatment periods, as we observed in a subset of mice and NHPs, and with no way to evaluate success or failure during or within months after the completion of a treatment regimen, this risk may not be worth taking.

A shortcoming of this study is that we did not directly compare the high dose/intermittent/long-term treatment to a more standard twice daily, 40-60 days BNZ dosing regimen that frequently fails in humans and other species. However, we have repeatedly made this comparison in nearly identical mouse studies [7] and a large number of studies have documented the inadequacies of this conventional daily dosing approach in primates and dogs [13, 14, 28, 29]. Furthermore, work herein and published elsewhere [24] in the De Brazza’s monkey demonstrates the superiority of the high dose, twice weekly BNZ protocol over both the standard daily regimen and a once-per-week protocol in the same animal. This latter study, as well as those in macaques herein, also show that even when they fail, these intermittent treatment protocols do not generate parasite population that are inherently resistance to BNZ. Thus, it is relatively safe and likely efficacious to ramp up the BNZ dosing regimen if a less intensive one initially fails, assuming that hosts can tolerate the increased dosing. Perhaps most importantly, we did not observe substantial toxicity in mice, dogs or NHPs when using these higher individual doses of BNZ over a longer period of time.

It should be emphasized that the dosing regimen evaluated here for BNZ may not necessarily apply to other anti-*T. cruzi* drugs. The unique mechanisms of action, cytotoxic/cytostatic activities, metabolism and tissue distributions of drugs combine to determine effective treatment regimens. We remain optimistic that shorter treatment courses may be possible using other compounds [20] although amastigote dormancy is likely to negate treatment periods of less than several weeks.

The pilot studies in dogs in this work provide the justification for more expansive trials that could further optimize the BNZ treatment protocol. In the southern U.S there are established enzootic cycles of *T. cruzi*, involving numerous triatomine vector species and a wide variety of mammalian hosts in addition to dogs and non-human primates [30, 31]. There are approximately >80 million dogs in U.S.[32] including many high value dogs (e.g. government and civilian (hunting) dogs) with extensive outdoor exposure and high opportunity for vector contact and acquiring the infection. In some settings, the new infection risk in such dogs can reach nearly 30% per year and these are frequently rapidly debilitating or fatal infections [3, 10, 33]. However, even in areas with more modest infection risk, the overall prevalence of *T. cruzi* infection in dogs in the southern U.S. ranges from 2 to >22% (reviewed in [34]). The apparent effectiveness of the BNZ protocol evaluated herein supports an effort to routinely diagnose and treat these infections and thus prevent morbidity and early mortality.

The suitability of the treatment protocol evaluated herein for use in humans is debatable as the total BNZ dose required for consistent cure is likely to be in the same range as that in the currently standard protocol. Thus, the high rate of adverse events with BNZ in humans may not be reduced using this intermittent protocol. However, until more effective and safer compounds are available, effective treatment regimens using existing drugs should be employed.

## Material and Methods

### Mice and infections

C57BL/6J mice (Stock No:000664) (C57BL/6 wild-type) were purchased from The Jackson Laboratory (Bar Harbor, ME). All the animals were maintained in the University of Georgia Animal Facility under specific pathogen-free conditions. *T. cruzi* tissue culture trypomastigotes of the wild-type Colombiana strain were maintained through passage in Vero cells (American Type Culture Collection (Manassas, VA)) cultured in RPMI 1640 medium with 10% fetal bovine serum at 37°C in an atmosphere of 5% CO_2_. Mice were infected, intraperitoneally with tissue culture trypomastigotes of *T. cruzi* and killed by carbon dioxide inhalation. This study was carried out in strict accordance with the Public Health Service Policy on Humane Care and Use of Laboratory Animals and Association for Assessment and Accreditation of Laboratory Animal Care accreditation guidelines. The protocol was approved by the University of Georgia Institutional Animal Care and Use Committee.

### BNZ treatment in mice

Infected mice were treated according to the indicated schedules. Benznidazole (BNZ – Elea Phoenix, Buenos Aires, Argentina) was prepared by pulverization of tablets followed by suspension in an aqueous solution of 1% sodium carboxymethylcellulose with 0.1% Tween 80 and delivered orally by gavage at a concentration dosage of 100 or 250 mg/kg body weight in 0.2 ml.

### T-cell phenotyping

Mouse peripheral blood was obtained by retro-orbital venipuncture, collected in sodium citrate solution, red blood cells lysed and the remaining cells were washed in staining buffer (2% BSA, 0.02% azide in PBS (PAB)) as previously described (Bustamante et al 2008, 2014, 2020). Whole blood was incubated with a major histocompatibility complex I (MHC I) tetramer containing the specific TSKB20 peptide (ANYKFTLV/Kb) labeled with BV421 (Tetramer Core Facility at Emory University, Atlanta, GA) and the following labeled antibodies: anti-CD8 FITC, anti-CD4 APC EF780, anti-CD127 PE (BD Bioscience, San Jose, CA). For whole blood, we lysed red blood cells in a hypotonic ammonium chloride solution after washing twice in PAB. We stained cells for 45 min at 4°C in the dark, washed them twice in PAB and fixed them in 2% formaldehyde. At least 500,000 cells were acquired using a CyAn ADP flow cytometer (Beckman Coulter, Hialeah, Florida) and analyzed with FlowJo software v10.6.1 (Treestar, Inc., Ashland, OR).

### Dog studies

Dogs for study were recruited from a network of kennels in central and south Texas with a history of triatomine vector occurrence and canine Chagas disease with owners who were willing to participate in the study. Informed consent was obtained from dog owners prior to their participation, and this study was approved by the Texas A&M University Institutional Committee on Animal Use and Care and the Clinical Research Review Committee (IACUC 2018-0460 CA).

At each sampling event, up to 6ml of blood was collected from dogs via venipuncture into clot activator and EDTA containing tubes which were shipped on ice overnight to the laboratory after which serum was separated and used for antibody tests, and the clot was used for DNA extractions and PCR testing [10]. The dog owners/kennel managers gave BNZ by placing the powdered drug in food. Dogs were owner-observed for physical and behavioral changes and a health check was carried out every 3-4 months throughout the duration of the study which included a physical examination, analysis of locomotion, postural reflexes and reaction to stimuli.

### Macaque Studies

The monkeys used in this study were 10 naturally infected, *Trypanosoma cruzi*-positive rhesus macaques (*Macaca mulatta*). Prior to being enrolled in the study, each macaque had been identified to be serologically positive for *T. cruzi* a minimum of 7 times and PCR-positive for *T. cruzi* a minimum of 2 times (Supplemental table 2 [35]). The animals were acquired from an approximately 1,000-animal breeding colony housed at the AAALAC-accredited, University of Texas MD Anderson Cancer Center, Michale E. Keeling Center for Comparative Medicine and Research (KCCMR) in Bastrop, Texas. The animals in this colony are specific-pathogen-free (SPF) for Macacine herpesvirus-1 (Herpes B), Simian retroviruses (SRV-1, SRV-2, SIV and STLV-1) and *Mycobacterium tuberculosis* complex. The macaque experiments were performed at the KCCMR and all protocols were approved by the MD Anderson Cancer Center’s IACUC and followed the NIH standards established by the Guide for the Care and Use of Laboratory Animals [35].

### BNZ treatment, health monitoring, and study protocol in macaques

This study was divided into two distinct phases. Treated animals in both phases of the study received a 37.5 mg/kg oral dose of powdered BNZ, hidden within preferred food treats. The control animals received similar treats at each study time point but without the addition of BNZ. The animals were isolated from their cagemates when presented with the treats and allowed time to consume the treat before being re-paired with their study cagemate. The vast majority of macaques consumed the treats without issue but where animals refused to eat the treat, a second and third attempt at oral-dosing the treats was undertaken as needed. Animals that refused to ingest the BNZ treats after three attempts were sedated and administered their BNZ dose through gastric gavage on the morning following the first attempted oral dosing. Through these combined efforts, all animals in this study received their appropriate BNZ dose within a 24-hour time period of when they were initially scheduled to receive the drug. All study animals were monitored at least twice a day by veterinarians and/or veterinary technicians. Baseline blood samples were collected as reference values for each animal and multiple blood collections were obtained throughout the study in order to ensure the overall health of the study animals. Physical exams were likewise performed whenever the animals were sedated for any blood draw during the study.

In the first phase of the study all 10 macaques were moved into study housing and allowed to acclimate while they were simultaneously trained for voluntary oral dosing over a one-month period. Following pre-study blood collections and health examinations, 8 of the 10 macaques (6 females and 2 males) began 1X per week 37.5 mg/kg BNZ dosing with the remaining two animals (both female) designated as untreated controls (Figure 3B). Other than animal T1-3 which had its BNZ treatment discontinued after 25 doses, the 1X per week dosing occurred over 28 weeks until dosing was stopped as a result of research restrictions associated with the Covid-19 pandemic. Health examinations, which included a physical examination, serum chemistry and CBC analysis, were performed at study week 2 and 4 for the eight treated macaques. Health examinations, with additional blood collections for study purposes, were also performed on study weeks 8, 16, and 24 for all 10 macaques. Following the initial cessation of BNZ dosing, four of the treated animals and one of the control animals then underwent a 10-week recovery period prior to being euthanized. Blood samples and fresh, frozen, and paraformaldehyde-fixed tissues were collected from these five animals. The remaining four treated animals and control animal underwent a 15-week recovery/washout period prior to starting the second phase of the study. Following the washout period, pre-study blood collections and health examinations were performed on all five animals. Four animals (2 female and 2 male animals) then began 2X per week 37.5 mg/kg BNZ dosing with the remaining animal (female) designated as an untreated control (Figure 3B). The 2X per week dosing was continued for 15 weeks (30 total BNZ treatments). Health examinations, which included a physical examination, serum chemistry and CBC analysis, were performed at week 2 and week 4 following the initiation of the 2X per week BNZ dosing for the four treated animals. Health examinations, with additional blood collections for study purposes, were also performed on study weeks 8 and 16 for all five animals. All five animals underwent a final health examination and then were euthanized following a four-week recovery period. Blood samples and fresh, frozen, and paraformaldehyde-fixed tissues were collected from the animals and processed for clinical tests and PCR detection of parasite DNA as previously described [20]

### De Brazza’s monkey

An 11 yr captive-bred male De Brazza’s monkey (*Cercopithecus neglectus*) in the Birmingham Zoo, Alabama, determined to be positive for *T. cruzi* infection based on serology, blood PCR and direct observation of parasites in blood, was submitted to multiple rounds of BNZ treatment, including a 35 mg/kg (5-fold the normal daily dose) once and then twice weekly, as indicated. A complete description of this case-study has been published elsewhere [24].

### Quantitative polymerase chain reaction in tissue samples and hemoculture

Mouse, dog or macaque tissue samples were processed for quantification of *T. cruzi* DNA by real-time polymerase chain reaction (qPCR) as previously described [7, 20]. For mice, approximately 100μl piece of tissue was minced finely using micro-dissecting scissors and DNA was extracted using the protocol included with the Qiagen DNeasy Blood and Tissue Kit (Qiagen, Hilden, Germany). For dog and macaque tissues a 8mm biopsy punch was employed to quickly collect both uniform and varied punches of approximately 100 μl from frozen tissues (Sklar instruments #96-1130). Single 100 μl tissue punches and a combination of five - 100 μl punches from different areas of tissue totaling 500 μl per pool. The spin-column protocol for “Purification of Total DNA from Animal Tissues”, (Milipore Sigma, Burligton, MA) was used as described by the manufacturer. All DNA samples were diluted to 25 ng/μl in nuclease free water. The generation of PCR standards and detection of parasite tissue load by qPCR was carried out as previously described [7, 8, 36]. The Biorad CFX manager software version 3.1 was used to analyze PCR data.

For hemocultures determinations, peripheral blood from mice, dogs or NHP was collected and cultured at 26°C in supplemented liver digest neutralized tryptose (LDNT) medium as described previously [37]. The presence of *T. cruzi* parasites was assessed every week for 3 months under inverted microscope.

### Multiplex serological analysis

Luminex-based multiplex serological assays were performed as previously described [10, 38, 39]. Recombinant *T. cruzi* proteins used included (TritrypDb.org identifiers: Tc1= fusion of TcBrA4_0116860 and TcYC6_0028190; Tc2= fusion of TcBrA4_0088420 and TcBrA4_0101960; Tc3 = fusion of TcBrA4_0104680 and TcBrA4_0101980; Tc4= fusion of TcBrA4_0028480 and TcBrA4_0088260; Tc5 = fusion of TcYC6_0100010 and TcBrA4_0074300; Tc7 = fusion of TcYC6_0083710 and TcBrA4_0130080; Tc8 = TcYC6_0037170; Tc11 = TcYC6_0124160; Tc17 = fusion of TcBrA4_0028230 and TcBrA4_0029760; Kn107 = TcCLB.508355.250; G10 = TcCLB.504199.20; Parvo = Recombinant Canine Parvovirus VP2 (MyBioSource.com) Tet = Tetanus toxoid (Sigma).

### Epimastigote growth inhibition

The assay of *in vitro* anti-*T. cruzi* activity on *T. cruzi* epimastigotes was performed based on the protocol described previously [40]. Approximately 25,000 log-phased epimastigotes from hemoculture-derived lines were subjected to serial 2-fold dilutions of BNZ for 72 hours. ATP production was measured as an indicator of viability using ATPlite™ Luminescence ATP Detection Assay System (PerkinElmer,). Luminescence was read using BioTek Synergy Hybrid Multi-Mode reader (BioTek,). The dose-response curve was generated by linear regression analysis with GraphPad Prism 5.0 (GraphPad Software *Inc*.). IC50 was determined as the drug concentration that was required to inhibit 50% of growth compared to that of parasites with no drug exposure.

### Statistical analysis

The non-parametric Mann-Whitney U tests, unpaired t-test and the mixed-effect analysis from the software GraphPad Prism v9.2.0 were used. Values are expressed as mean ± SEM.

## Acknowledgements

We thank the kennel owners and dog managers, Keswick Killits for assisting with BNZ powder preparation, and Drs. Erin Edwards at Texas A&M Veterinary Medical Diagnostic Laboratory for assistance with the study. Insud/VetPharma, the Mundo Sano Foundation and Humanigen donated benznidazole for the study. Facilities in the CTEGD Cytometry Shared Resource Lab were critical to the study. Funding for the study came from National Institutes of Health grants R01 AI151148 and R01 AI125738 to RLT, NIH grant R21 AI142469 to JMB and RLT, Veterinary Pharmacology Research Foundation/American Veterinary Medical Foundation Research Grant; University of Texas Southwestern / Texas A&M University Pilot Award for the NIH Clinical Translational Science Award (CTSA) 1UL1TR003163-01A1; The Harry Willett Foundation.

